# Analysis of genotype by environment interactions in a maize mapping population

**DOI:** 10.1101/2021.07.21.453280

**Authors:** Asher I. Hudson, Sarah G. Odell, Pierre Dubreuil, Marie-Helene Tixier, Sébastien Praud, Daniel E. Runcie, Jeffrey Ross-Ibarra

**Affiliations:** Department of Evolution and Ecology, University of California, Davis; Center for Population Biology, University of California, Davis; Department of Plant Sciences, University of California, Davis; Center of Research of Chappes, Limagrain, route d’Ennezat, 63720, France; Genome Center, University of California, Davis

## Abstract

Genotype by environment interactions are a significant challenge for crop breeding as well as being important for understanding the genetic basis of environmental adaptation. In this study, we analyzed genotype by environment interaction in a maize multi-parent advanced generation intercross population grown across five environments. We found that genotype by environment interactions contributed as much as genotypic effects to the variation in some agronomically important traits. In order to understand how genetic correlations between traits change across environments, we estimated the genetic variance-covariance matrix in each environment. Changes in genetic covariances between traits across environments were common, even among traits that show low genotype by environment variance. We also performed a genome-wide association study to identify markers associated with genotype by environment interactions but found only a small number of significantly associated markers, possibly due to the highly polygenic nature of genotype by environment interactions in this population.

## Introduction

Both the effect of a given genotype on a trait, and the impact of that effect on fitness, often vary across environments. Such genotype by environment interactions (GxE) are widespread, and have been commonly observed in plants (Des Marais *et al*. 2013). GxE interaction is of interest for multiple reasons: it provides insight into the physiological processes and genetic architecture underlying individual traits, is likely crucial for local adaptation of populations to different environments, but may also limit the response to selection (Allard and Bradshaw 1964; Kawecki and Ebert 2004).

While alleles affecting a trait will demonstrate GxE for fitness across environments when there is selection for different trait optima, it is also often observed that the effect of individual alleles on traits will vary as well. This indicates that these alleles affect plasticity and they may be present in a population due to selection for or against plasticity (Josephs 2018). Alternatively, they may be deleterious but rarely exposed to environments in which they are selected against, or unassociated with fitness and selectively neutral (Des Marais *et al*. 2013; Paaby and Rockman 2014).

One avenue to study GxE is to search for individual loci with changing effects on traits or fitness across environments. Multiple studies have identified loci that contribute to GxE (several of which are reviewed in Josephs (2018)). Loci which contribute to GxE include the Eda locus in threespine stickleback fish, which is associated with adaptation to the freshwater environment, and Sub1A in rice, which is associated with tolerance to submergence (Barrett *et al*. 2008; Xu *et al*. 2006). Genome-wide association studies (GWAS) have also been used to identify alleles significantly associated with GxE, including shade response and drought response in *Arabidopsis thaliana* (Filiault and Maloof 2012; El-Soda *et al*. 2015).

Individual traits do not exist in a vacuum, however, and alleles that affect one trait often have pleiotropic effects on others. Indeed, the outcome of selection on a trait depends crucially on the genetic variance-covariance matrix (G-matrix), which describes how the genetic value at one trait covaries with genetic values at other traits (Lande 1979). Genetic covariation between traits can have profound impacts on the genetic response to selection, either hindering or facilitating trait response. For example, genetic covariance between traits that are both associated with fitness can lead to trade-offs between those traits if the covariances with fitness are mismatched.

But the G-matrix itself is not constant, as GxE at underlying loci may impact trait variation and covariation among traits (Wood and Brodie 2015). If in a different environment the covariance of a trait with fitness or other traits is weakened or changes sign, it may indicate that the selection or trade-off does not exist in the new environment (Sgr’o and Hoffmann 2004). As GxE contributes to the G-matrix within each environment, understanding the G-matrix in multiple environments may illuminate the causes of GxE. If the genetic covariance between two traits changes between environments and GxE is observed, then a change in the pleiotropy of the underlying loci may be responsible for both the changes in the genetic covariance and GxE.

Maize is a crop species adapted to a wide diversity of environments, from temperate to tropical and from low to high altitude (Hake and Ross-Ibarra 2015). GxE has been shown to be an important contributor to many traits in maize, including grain yield (Gage *et al*. 2017; Gates *et al*. 2019; Rogers *et al*. 2021). Nonetheless, identification of GxE in maize, as in many species, is complicated by issues of population structure and the low minor allele frequency of most polymorphisms (Korte and Farlow 2013). To circumvent these issues, we investigated the genetic basis of GxE in maize in a multiparent advanced generation intercross (MAGIC) population of 16 diverse temperate maize lines (Odell *et al*. 2021). We grew the MAGIC hybrids across five contrasting temperate environments with diverse management practices in order to capture a broad range of GxE relevant to the conditions the parental lines would be grown in.

We find that GxE contributes as much as genotypic main effects to variance for some traits. While GxE interactions are significant, genome-wide association only finds a small number of markers significantly associated with GxE interactions, perhaps reflecting the highly polygenic nature of most traits. Nonetheless, estimation of the G-matrix in each environment reveals that changes in genetic covariance are common and may be contributing to observed GxE. For example, we find that while only a small proportion of variance in flowering time depends on GxE, the genetic covariance between flowering time and grain yield is strongly affected by the environment.

## Materials and Methods

### Plant materials

We developed a maize multi parent advanced generation intercross (MAGIC) population by repeatedly crossing the offspring of sixteen maize inbred lines to generate recombinant individuals (Odell *et al*. 2021). After eight generations of intercrossing, we generated a population of 344 doubled haploids (DH) lines. DH lines were crossed to an inbred tester to make F1 plants.

### Phenotype Data

The MAGIC F1 plants were phenotyped in four different field locations in four different years, resulting in five distinct environment-years (Figure S1, Table S1). The environment-years included Blois, France in 2014 and 2017, Nerac, France in 2016, St. Paul, France in 2017, and Graneros, Chile in 2015. In each environment-year, two plots of around 80 plants were grown for each genotype. The fields in environment-years Blois 2014, Blois 2017, and Graneros 2015 all received consistent irrigation. The field in Nerac 2016 was not actively irrigated from vegetative phase through flowering, causing drought stress through most of the life cycle. The field in St. Paul 2017 was not irrigated during vegetative phase but was irrigated during flowering to allow plants to recover from the earlier drought stress. We measured the following traits: male flowering date, female flowering date, anthesis-silking interval (ASI), plant height, percent harvest grain moisture (HGM), grain yield, and thousand kernel weight (TKW) (adjusted to 15% humidity), where values were averaged over plots. Both flowering time phenotypes were measured as the sum of degree days since sowing with a base temperature of 6^*°*^C (48^*°*^F). Male flowering date was considered as the growing degree days until 50% of plants in a plot were shedding pollen on approximately one quarter of the central tassel spike. Data was also collected from an additional environment, Szeged, Hungary in 2017. We did not use this data in the analyses presented here as flowering date was not collected on the same schedule as in the other environments and this caused issues with the GxE analyses. Data from Szeged is available in the data repository associated with this paper. Between 292 and 309 of the MAGIC F1 lines were grown in each environment. There were a total of 325 lines that had both genotype data and phenotype data from at least one environment. For each of these lines we calculated best linear unbiased predictor (BLUP) scores for all seven phenotypes, combining measurements from all environments to get estimates of the genetic contribution to the phenotype for each MAGIC line (Aulchenko *et al*. 2007).

### Genotyping

We genotyped each of the DH lines using the Affymetrix^®^ Axiom^®^ Maize Genotyping Array, which successfully genotyped 551,460 SNPs. The probability of each founder contributing to each segment in the genome was imputed from the genotyped SNPs (Odell *et al*. 2021).

### Statistical analysis

All statistical analyses were performed with R (R Core Team 2020). Plots were made with *ggplot2* (Wickham 2016).

### Estimating kinship

Kinship matrices for the DH lines were estimated from the genotyped SNPs using the VanRaden method as implemented in the R package *sommer* (Covarrubias-Pazaran 2016; VanRaden 2008). SNPs were first filtered for linkage disequilibrium using Plink (Purcell *et al*. 2007). In order to perform genome-wide association analyses, we used the leave one chromosome out method (Lippert *et al*. 2011).

### Genotype x environment interactions

Variance components for each trait were estimated using the R package *sommer*. We used the formula:

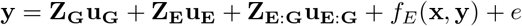

Where **y** is a vector of *n* observations from individual plots of a single trait including both plots of all lines in all environments, **Z**_**G**_ is a *n* × *r* design matrix for the genotypic main effects of the *r* lines, **Z**_**E**_ is a *n* × 5 design matrix for the environmental main effect, **Z**_**E**:**G**_ is a *n* × 5*r* design matrix for genotype specific effects in each environment, **u**_**G**_ is a length *r* vector of random genotypic effects, **u**_**E**_ is a length 5 vector of environmental random effects, **u**_**E**:**G**_ is a length 5*r* vector of random GxE effects with same variance and covariance among environments, *f*_*E*_(**x, y**) is a two dimensional spline for the effect of the x/y position in the field nested within environment, and **e** is the error.

### GWAS

Genome-wide association analyses for loci contributing to GxE interactions were performed with the R package *GridLMM* (Runcie and Crawford 2019). Imputed founder probabilities at each locus were used as markers, meaning that at each marker we asked if the identity of the founder which contributed that genomic region at a given locus was a significant predictor of differences in plasticity among the hybrids.

We modeled GxE in three ways:

i. Main effect across environments and deviation effect within environments We tested whether a locus had a different effect on a trait in two environments: Blois 2017 and Nerac 2016. We chose these two environments because they were respectively the highest and lowest yielding environments. The model for this GWA was:

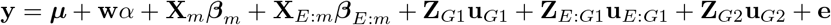

Where **y** is a vector of *n* observations from individual plots of a single trait including both plots of all lines in all environments, ***µ*** is a constant length *n* vector of the average trait value across the two environments, **w** is a length *n* design matrix of environmental effects taking values of -1 and +1 according to the environment (1 for Blois 2017 and -1 for Nerac 2016), *α* is a scalar representing 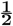 the deviation of trait means between the two environments. **X**_*m*_ is a *n* × 16 matrix, where the *k*th column is the probability that each of the *n* individuals inherited from the *k*th founder at marker *m*, **X**_**E**:**m**_ is an *n* x 16 matrix formed by multiplying **w** with each column of **X**_*m*_, ***β****m* is a vector of main effects of the founder alleles averaged over the two environments, ***β****E*:*m* is a vector of differences between the founder allele effects between the two environments, **Z**_*G*1_ is a *n* × *r* design matrix of additive genotypic effects, **Z**_*E*:*G*1_ is a *n* × *r* design matrix of genotype deviations formed by multiplying each column of **Z**_*G*1_ by **w, Z**_*G*2_ is a *n* × *r* design matrix of non-additive genotypic effects, **u**_*G*1_ is a vector of additive genotypic effects averaged over the two environments, **u**_*E*:*G*1_ is a vector of additive genotypic deviations between the two environments, **u**_*G*2_ is a vector of non-additive genotypic effects averaged across the two environments, and **e** is a vector of error terms. **u**_*G*1_ and **u**_*E*:*G*1_ both have covariance proportional to **K**, where **K** is the additive genetic relatedness matrix, and **u**_*G*2_ and **e** both have covariance proportional to the identity matrix. The statistical test to identify markers influencing GxE was against H0: ***β****E*:*m* = **0**.
ii. Plasticity We tested whether a locus had an effect on the slope of the observations of a genotype across the mean phenotypic value of all genotypes in an environment. The model is the same as in i) except for the following: we now include all 5 environments, **w** is a length *n* vector with each element taking the mean value of the phenotype within the environment of the observation, and ***µ*** is a length *n* vector of the mean value of the phenotype within the environment of the observation.
iii. Finlay-Wilkinson GWAS Finally, we tested whether a locus had an effect on the slope of the observations of a genotype across the mean grain yield of all genotypes in an environment. Mean grain yield here serves as a proxy for stress or environment quality and as such this GWA is testing whether a locus affects the response to stress. This is known as a Finlay-Wilkinson analysis (Finlay and Wilkinson 1963). For this analysis, a quantile plot of p-values indicated that the test was poorly calibrated. Instead of asking whether allowing a marker to have a slope across environments improved prediction of a trait in each environment as in (ii), we thus asked whether the marker significantly predicted the slope of each genotype.

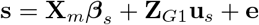

Where **s** is a length *r* vector of slopes for each genotype of trait values on mean grain yield in each environment, ***β****s* is a vector of marker effects, and **u**_*s*_ is a vector of genotypic effects with covariance proportional to **K**. Other model terms are as in (i).

To determine significance thresholds for the first two models, we permuted phenotypic values among lines within each environment and ran the GWA 100 times. For the third model, we permuted the slopes among the genotypes and ran the GWA 100 times.

### The G-matrix across environments

We estimated the G-matrix in each environment using the R package *brms* (Bürkner 2017). brms implements Bayesian multilevel models using Markov chain Monte Carlo (MCMC) algorithms. This is important as the samples from the MCMC chains allow us to estimate uncertainty and significance in our downstream analyses. We used the model:

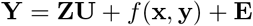

Where **Y** = [**y**_**1**_…**y**_**5**_] and **y**_**i**_ is a vector of *n* observations for the *i*th trait, **Z** is a *n* × *r* design matrix of genotypes, **U** and **E** are random effects drawn from matrix normal distributions: **U** ∼ *MN*_*r*,5_(0; **I**_**r**_, **G**), **E** ∼ *MN*_*n*,5_(0; **I**_**n**_, **R**) and **I**_**r**_ is the *r* × *r* identity matrix where *r* is the number of lines grown in an environment, **I**_**n**_ is the *n* x *n* identity matrix where *n* is the number of observations, and **G** and **R** are 5 × 5 genetic variance-covariance and residual variance-covariance matrices estimated from the data. *G* and *R* are parameterized as the products of standard deviations and correlation matrices with a half Student-T distribution and LKJ-correlation prior. *f* (**x, y**) is a two dimensional spline for the effect of the x/y position in the field. The standard deviations of the two splines have half Student-T distributions as priors.

All traits were scaled by the mean and centered before analysis in order to make them unitless and improve model convergence, with the exception of ASI which was scaled by the mean of female flowering as those were on the same measurement scale. The G-matrices we estimated were broad sense G-matrices as they included both additive and non-additive sources of genetic variance. We assessed convergence by checking that all statistics output by brms — such as 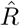 and the number of divergent transitions — were within recommended ranges and by visually inspecting the trace and autocorrelation of model parameters. Covariances were considered significant if the interval spanned by the 2.5% and 97.5% quantiles of the posterior samples did not contain zero. To determine whether the covariance between two traits differed significantly between environments, we found the difference between the MCMC samples for the two environments and determined whether the interval spanned by the 2.5% and 97.5% quantiles of the differences overlapped zero.

To quantitatively assess differences among the G-matrices estimated in the five environments, we performed eigenanalysis of a covariance tensor as described in Aguirre et al. (Aguirre *et al*. 2014). The tensor approach is a geometric approach founded on the diagonalization of symmetric matrices, and is mainly used to calculate a set of orthogonal axes known as eigentensors that describe coordinated changes in the elements of the original matrices being compared. Eigentensors describe which elements of a set of matrices most contribute to variation among those matrices. Eigentensor analysis was performed on the posterior median G-matrices. Uncertainty in the eigentensors was estimated by performing eigentensor analysis on the MCMC samples of the G-matrices. Finally, to determine whether an eigentensor explained more of the variation among G-matrices than would be expected by chance, we shuffled the real phenotypic data among environments, estimated G-matrices, and asked whether the eigentensors of the randomized G-matrices explained as much of the variation as the MCMC samples from the real data.

## Data availability

Phenotypic and environmental data are located on Figshare at (available on publication). Genotypic data will be made available through a data repository associated with companion paper (Odell *et al*. 2021) (available on publication).

## Results

We evaluated 7 phenotypes for each of 344 doubled haploid (DH) lines in replicated trials across 5 environments that varied in temperature, daylength, and watering or drought conditions. Each DH line was genotyped for 551,460 SNPs, allowing us to identify ancestry segments along the genome.

### Genotype x environment interactions

Genotypic main effects and GxE interactions contributed a significant amount of the variance of all measured traits (Figure 1). Across environments, it was common for the rank of DH lines for grain yield to change, indicating that individual lines were generally not high yielding in all conditions (Figure 1A). Anthesis-silking interval (ASI) showed a qualitatively similar pattern of rank-changing, while some traits such as thousand kernel weight (TKW) showed less dramatic GxE (Figure S2). The proportion of variance due to main genotypic effects ranged from 0.34 for grain yield to 0.72 for male flowering date (Figure 1B). For grain yield and HGM, GxE interactions contributed an amount of variance similar to the amount contributed by genotypic effects. For flowering time, TKW, and plant height, GxE interactions contributed less of the variance than main genotypic effects.

**Figure 1.**
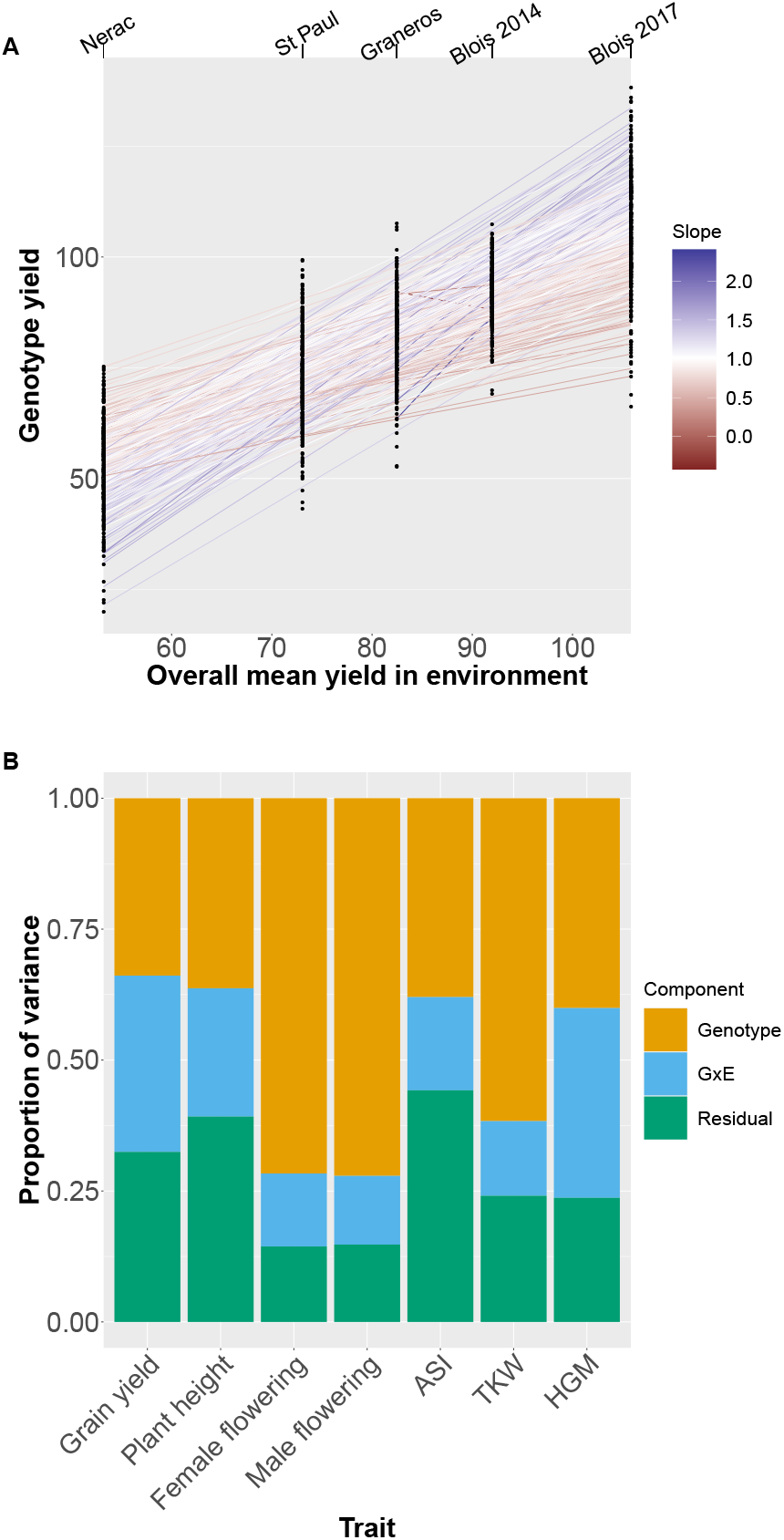
A) Mean yield of all genotypes in each environment. On the X axis environments are plotted by the mean yield across all genotypes in that environment. Points are mean yields of individual genotypes. Lines are the slope of a genotype’s mean yield in each environment on the mean yield of all genotypes in that environment. The color of the line corresponds to the slope; a slope greater (or less) than one indicates a genotype more (or less) responsive to the environment than average. B) Restricted maximum likelihood estimates of variance components for each trait across all environments.

### GWAS

Our test of the deviation effect of a marker within environments did not recover any markers significant at the 5% permutation threshold for any trait. In contrast, our plasticity GWAS identified two peaks which were significant at the 5% significance level, which were for ASI and female flowering (Figures 2A, S3). Neither of these peaks overlapped with QTL peaks for main effects in this population (Odell *et al*. 2021). The peak for ASI on chromosome 1 appears to be driven by the effect of the FV2 founder, which has a small effect in environments where ASI is close to zero but strongly increases the magnitude of ASI in environments where average ASI is greater (Figure 2B). Patterns of identity by descent at the genomic region surrounding the peak identified unique haplotypes for 15 of the founders (Odell *et al*. 2021), but a PCA of the SNPs in the region did not indicate that the FV2 haplotype was strongly diverged from other founders (Figure S4). The peak for female flowering on chromosome 4 appears to be driven by founder A654, but the marker effects for this founder appeared unrealistically strong and likely reflect an artifact of the extremely low sampling of this founder among the DH lines. In addition to these two associations at the 5% level, we detected one peak which was significant at the 10% level for grain yield (Figure S5). Our Finlay-Wilkinson GWAS uncovered one peak significant at the 5% level for ASI (Figure S6). However, the founder whose effect appears to be driving this peak also appears to be underrepresented at this locus and only one line has a greater than 0.8 probability of carrying this founder allele. As a result, this peak is likely to be a statistical artifact.

**Figure 2.**
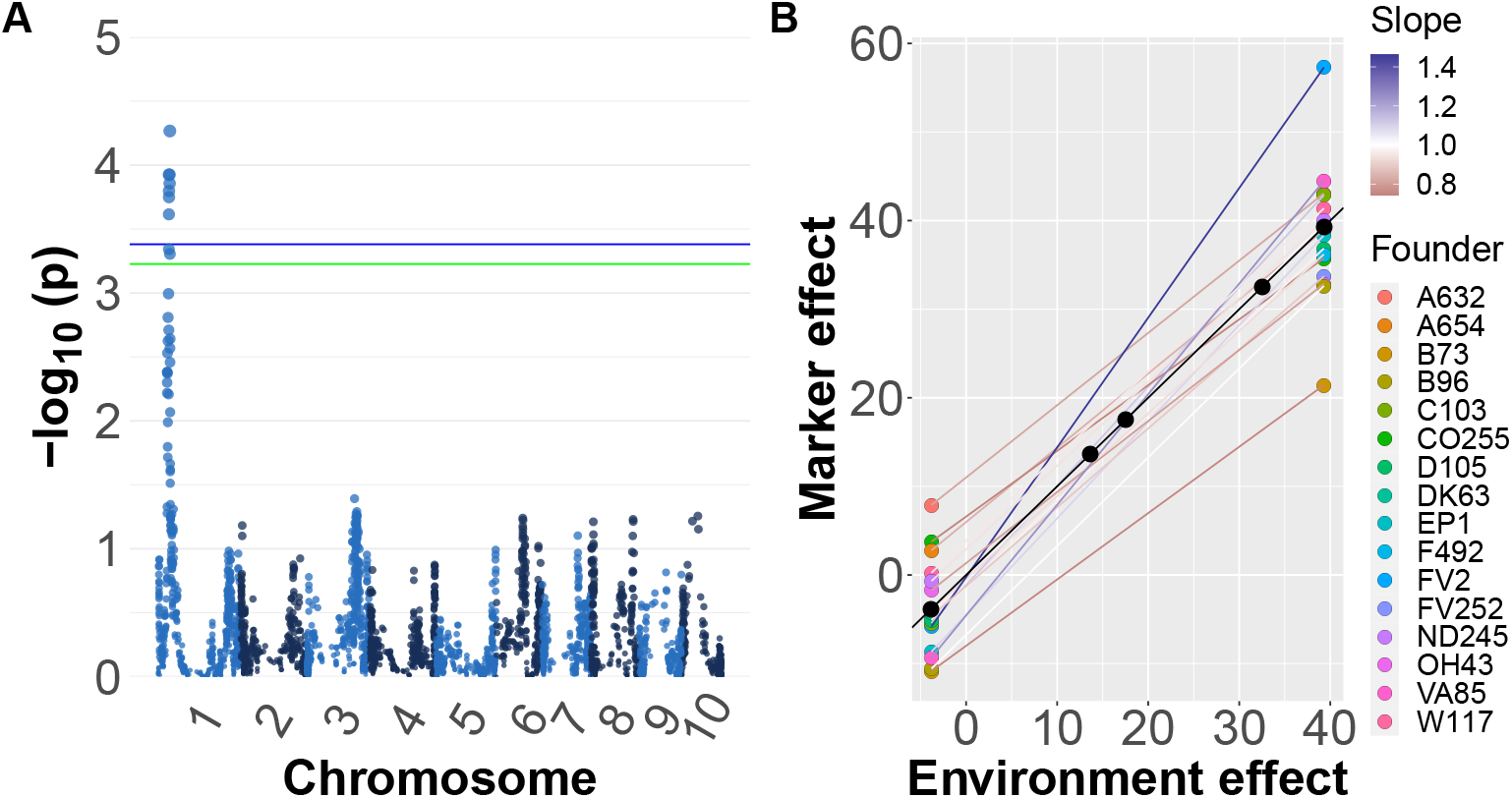
A) Manhattan plot for plasticity (model ii) GWAS on ASI. The blue and green lines represent the 5% and 10% significance levels based on permutation tests, respectively. B) Estimated effect of founder ancestry on plasticity for the most significant marker. The slope of a line indicates the plasticity of that haplotype and the difference in slopes is GxE. The color of the line corresponds to the slope; a slope greater (or less) than one indicates a genotype more (or less) responsive to the environment than average.

### The G-matrix across environments

To understand how the environment affected pleiotropy, we estimated the genetic variance/covariance matrix (G-matrix) of five traits in each environment (Figures 3A, B, S7). We dropped male flowering date and HGM from this analysis because models including those traits failed to converge; in analyses run on subsets of these traits we found that male flowering date was highly correlated with female flowering date and HGM had very low covariance with the other traits. Comparisons of the 95% credible intervals of the difference between individual genetic correlations revealed numerous differences among environments (Figure S8). Both the genetic variances of individual traits and the covariances between traits differed across environments (Figure 3A, B). As the traits were mean scaled, the variances presented in Figure 3A are not heritabilites, which is the genetic variance scaled by the phenotypic variance. Importantly, mean-scaled genetic variances are not affected by the amount of residual variance, which means that a trait with high genetic variance relative to the mean along with high environmental variance can have low heritability but high mean-scaled genetic variance. (Houle 1992). We found that grain yield generally had high mean-scaled genetic variance in each environment, and the single highest mean-scaled genetic variance of any trait in any environment was grain yield in Blois 2017. In one case, the sign of a genetic covariance changed: the genetic covariance between grain yield and female flowering date was positive across all environments except in Nerac 2016. The median posterior values of some other genetic covariances also switched signs between environments, but based on credible intervals we cannot state that they switched with confidence.

**Figure 3.**
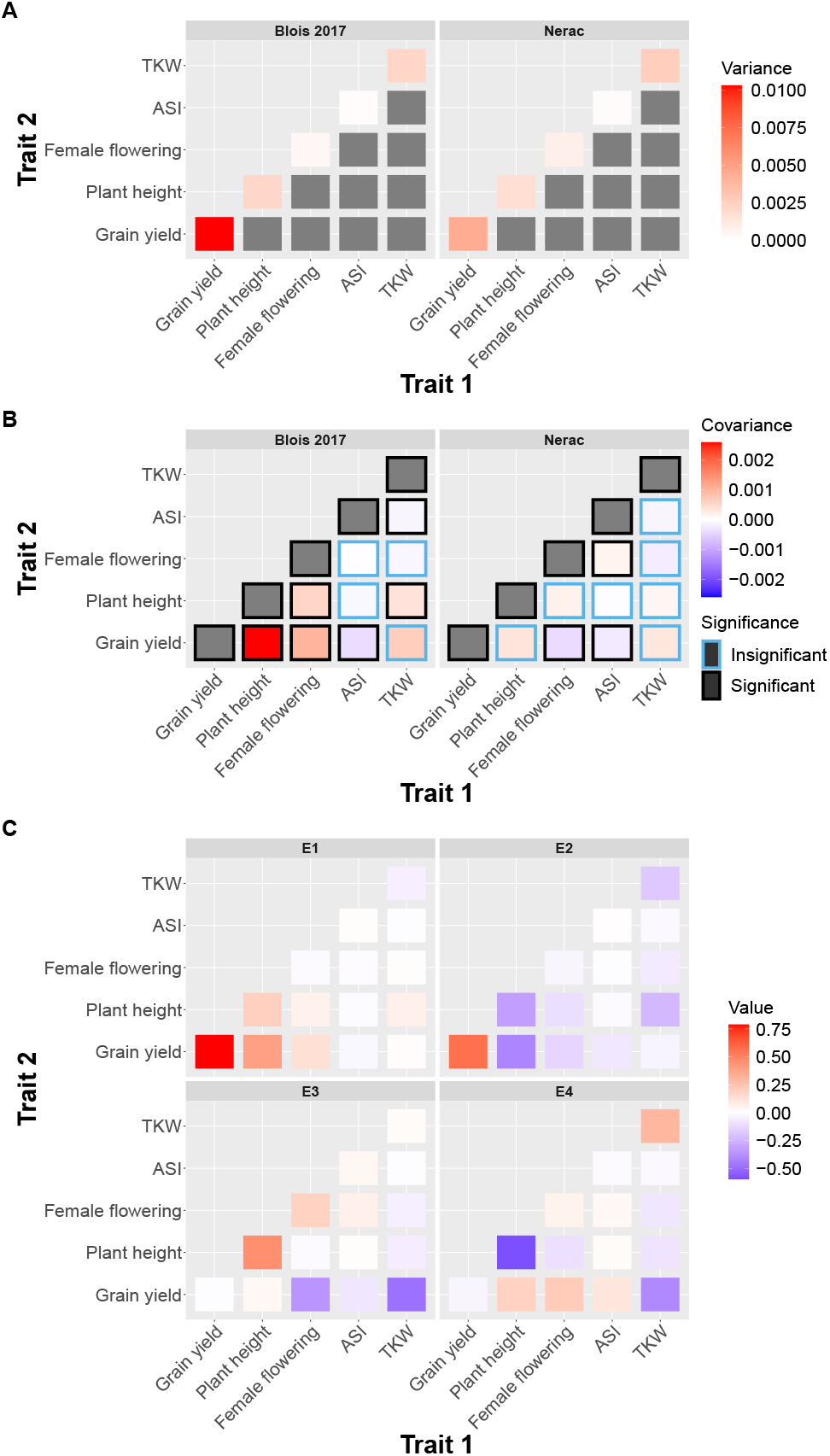
A) The genetic variances and covariances of the highest yielding environment (Blois 2017) and the lowest yielding environment (Nerac 2016). Genetic variances are on the upper row while genetic covariances are on the lower row. Traits are mean scaled. If a covariance is surrounded by a black frame, the value is significantly different from zero. Note that the scales on the upper and lower rows are different. B) Contributions of elements in the genetic variance-covariance matrices to the first four eigentensors of the set of genetic variance-covariance matrices. Elements on the diagonal are genetic variances of traits and elements on the off-diagonals are genetic covariances between traits. The color of a square represents the strength of the contribution of that element to the eigentensor, which is not dependent on the sign.

To quantitatively assess how individual elements of the G-matrix contributed to variation among environments, we performed an eigentensor analysis. The eigentensors of a set of G-matrices describe independent dimensions of variation among the G-matrices and can be used to identify which elements are contributing the most variation among the set. All of the four nonzero eigentensors explained significantly more variance than expected from permutations (Figure S9). The element of the G-matrix that most contributed to the first eigentensor was genetic variance for grain yield (Figure 3C). When plotting each environment on this eigentensor, Blois 2017 is strongly differentiated from the other environments, which is probably due to the genetic variance for grain yield being the highest in this environment (Figure S10). The genetic variance for grain yield also contributed strongly to the second eigentensor, while the genetic covariance between plant height and grain yield and the genetic variance of plant height contributed in the opposite direction. The third eigentensor described a contrast between genetic variance for plant height on the one hand and the genetic covariances between both female flowering date and TKW with grain yield on the other. Nerac is strongly differentiated on this eigentensor. While the covariance between female flowering and grain yield is not the only element of the G-matrix contributing to the third eigentensor, it is worth noting that Nerac is the only environment in which this covariance is negative.

## Discussion

### Genotype x environment interactions

Genotype x environment interactions are known to be important for many agronomically important traits in maize, and our results on the relative importance of GxE across traits confirm these earlier findings. For example, male and female flowering date have been shown to be influenced predominantly by additive genetic effects and are not strongly influenced by GxE interactions (Buckler *et al*. 2009; Rogers *et al*. 2021), while grain yield and HGM have large GxE variance components relative to main genotype effects (Gage *et al*. 2017; Rogers *et al*. 2021). We find similar results in our analysis, indicating that this may be a consistent pattern for diverse maize germplasm in temperate environments.

If genotypes are adapted to different environments, we would expect to see GxE for fitness related traits. The high variance contributed by GxE to grain yield seen in this study thus indicates that the founder maize lines, despite all having been bred in temperate environments, still carry many alleles that are differentially adapted to this set of environments. For traits that are further removed from fitness it is less clear how to interpret the contribution of GxE. It may be that the GxE we observe for a trait like HGM, which has a high proportion of GxE variance and a low genetic covariance with grain yield, is an example of neutral plasticity and is not under strong selection (Des Marais *et al*. 2013).

### GWAS

Despite the presence of substantial GxE variance for several traits, we found relatively few markers which were significantly associated with GxE. One possible explanation is that the GxE variance we observed is largely polygenic and caused by many loci of small effect which we did not have power to detect with our GWAS. Previous studies investigating loci with main effects on traits such as grain yield and flowering time in maize have found that they are highly polygenic (Buckler *et al*. 2009; Dell’Acqua *et al*. 2015). It may not be surprising then if GxE for these traits also has a similarly polygenic basis. Grain yield is a highly integrated trait dependent on the interaction of many other traits with the environment; if those traits have a complex basis and different optima within different environments, then it would not be surprising to observe large GxE variance at the level of genotype while not observing significant GxE effects for individual loci.

### The G-matrix across environments

The G-matrix has previously been shown to differ as much between environments as between populations (evidence reviewed in (Wood and Brodie 2015)). Our work shows that the G-matrix differs across environments in a multiparent population of temperate maize lines. We find that these differences include both changes in the magnitude of genetic variances and covariances as well as changes in the sign of genetic covariances. The highest mean-scaled genetic variance we observed was for grain yield in Blois 2017, and in general grain yield had high mean-scaled genetic variance compared to other traits within each environment. This is in contrast to the finding that grain yield had the lowest heritability across all environments. This finding fits with previous work finding that fitness proximal traits frequently have low heritability but high mean-scaled genetic variance, possibly because of high residual variance for fitness proximal traits reducing heritability (Houle 1992).

The magnitude of the genetic covariances between traits can be reduced solely as a function of reduced genetic variance for one or both of these traits without a change in the relationship between them. However, by looking at genetic correlations, we show that the correlations between traits varied across environments beyond effects of the differences in the variances (Figure S8). Additionally, changes in the genetic variance alone will not cause the covariance between traits to change sign, which we also see for some combinations of traits. Particularly striking was the change in sign for the genetic covariance between grain yield and female flowering date observed in the most stressful environment, Nerac 2016. This environment was the only one in which the genetic covariance between grain yield and female flowering date was negative. Previous work has shown that flowering time is important for adaptation to drought stress (reviewed in (Kazan and Lyons 2016)). Nerac 2016 experienced a drought from vegetative growth through maturity. Early flowering in this environment was genetically correlated with higher yields, suggesting that early flowering may have been a means to escape drought stress. The change in sign of the covariance is noteworthy given that we observed low GxE variance and high genotypic variance for female flowering date while simultaneously observing high levels of GxE variance for grain yield. This indicates that genotypes were relatively consistent in their flowering time across environments but that late flowering genotypes were higher yielding in most environments and lower yielding in one environment. In this way, a change in the genetic covariance between two traits (grain yield and female flowering) across environments may be contributing to GxE in one of those traits (grain yield), and provides an illustrative example of how traits that themselves show little GxE may nonetheless contribute to GxE for fitness.

While differences between environments presumably shape these changes in the G-matrix, previous work has found that neither measures of environmental novelty nor differences in phenotypic means shaped the G-matrix when looking across all the studies in a meta-analysis (Wood and Brodie 2015). In our analysis we find a similar result; differences between the G-matrices estimated in each environment are largely idiosyncratic and do not correspond with levels of stress or water availability. Eigentensor analysis reveals that each of the main directions of variation across G-matrices correspond mostly to the differentiation of one or at most two of the environmental G-matrices from the others. Previous work investigating the G-matrix of plant populations grown in well-watered and drought environments has been inconsistent in terms of whether drought stress increases or decreases genetic variance and how it affects the genetic correlation between flowering time and yield (Manzaneda *et al*. 2015; Sherrard *et al*. 2009).

Additionally, both the severity and timing of drought seem to be important in determining the effects of water deficit on covariances between traits. In this study we find that in Nerac, the most drought stressed environment, the genetic covariance between flowering time and yield is negative and that this genetic covariance contributes to differentiating it from the other environments. The fact that the genetic covariance between flowering date and grain yield in the other water deficit environment, St. Paul, was not significantly negative may be because that population was given water during flowering while in Nerac water deficit extended through flowering. It appears that how the G-matrix is affected by environmental stress is highly dependent on the species and population studied and the exact stress applied.

## Conclusion

Using a MAGIC population of maize grown in five environment x year combinations we were able to analyze the genetic basis of GxE in a set of diverse maize lines. We observed GxE variance for all traits and for some traits we observed comparable amounts of genotypic and GxE variance. Estimating the G-matrix within each environment revealed that changes in genetic variances and covariances across environments were common. Notably, the genetic covariance between yield and female flowering time was positive in most environments but negative in one of the environments. GWAS identified one locus significantly associated with GxE for anthesis-silking interval. Given the substantial GxE variance, the low number of significant loci suggests that GxE for the traits we analyzed may have a polygenic basis.

## Acknowledgments

We want to thank Silas Tittes and Sarah Turner-Hissong for their advice on statistical analysis. A.I.H. was supported by National Science Foundation Graduate Research Fellowship 1650042. J.R.I. was supported by USDA Hatch project CA-D-PLS-2066-H 548. S.O. was supported by the UC Davis Dept. of Plant Sciences and NSF grant 1754098. D.E.R. was supported by USDA Hatch project 1010469 and USDA NIFA grant no. 2020-67013-30904.

## Supplemental figures

**Table S1.** Features of the five growing environments.

| Environment-Year | Mean temperature (°C) | Mean relative humidity (%) | Mean precipitation (mm) | Water treatment<sup>a</sup> |
| --- | --- | --- | --- | --- |
| Blois 2014 | 16.7 | 75.2 | 2.19 | OPT |
| Blois 2017 | 17.0 | 72.3 | 1.71 | OPT |
| Graneros 2015 | 20.1 | 55.1 | 0.266 | OPT |
| Nerac 2016 | 19.1 | 74.9 | 1.15 | Early term |
| St. Paul 2017 | 20.3 | 65.4 | 1.12 | Recovery |

<sup>a</sup>OPT is optimum watering, Early Term is water deficit during vegetative growth through maturity, and Recovery is water deficit during vegetative growth with recovery at flowering time.

**Figure S1.**
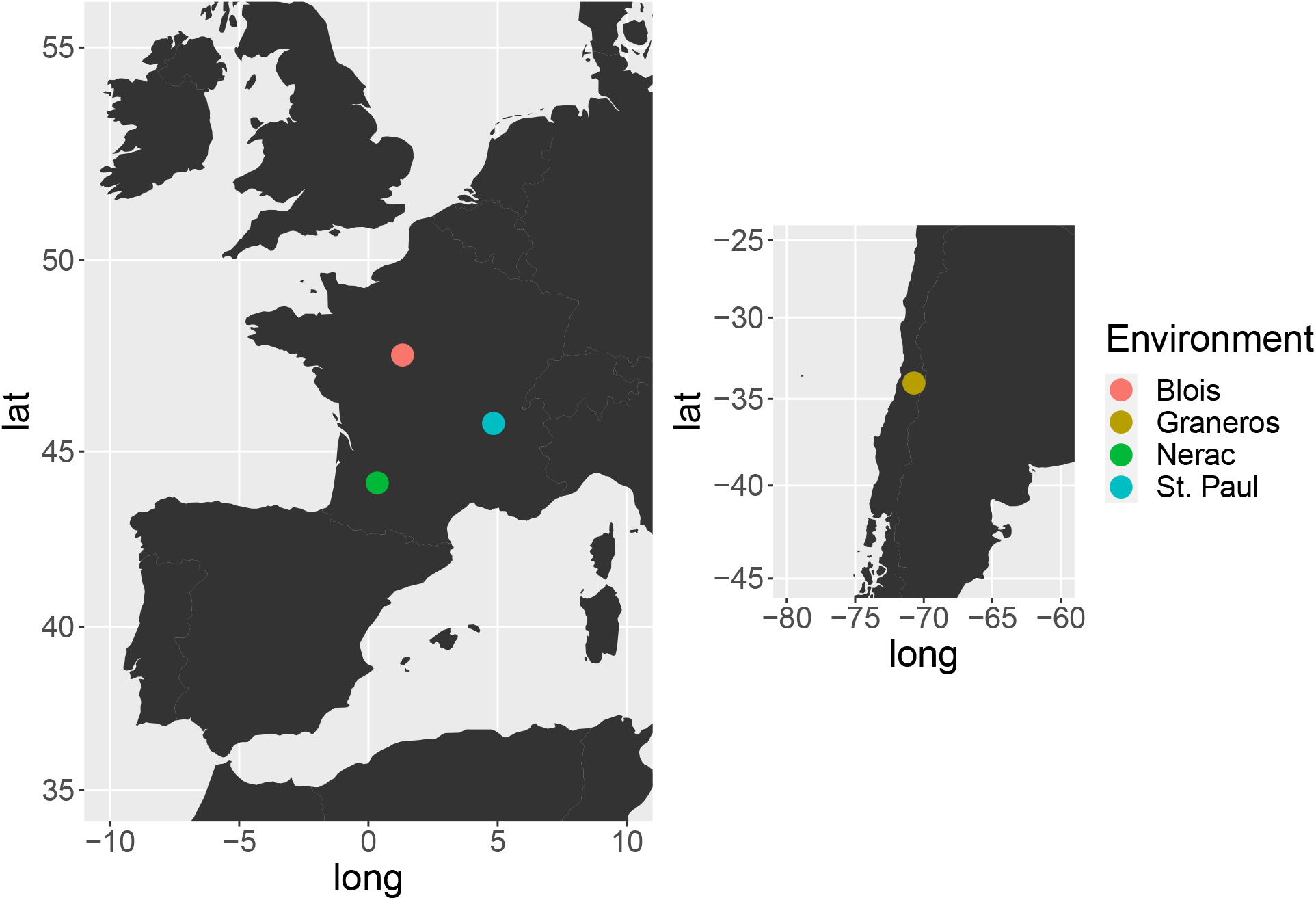
Locations of the environments the MAGIC population was grown in. In one environment (Blois) the MAGIC population was grown in two years.

**Figure S2.**
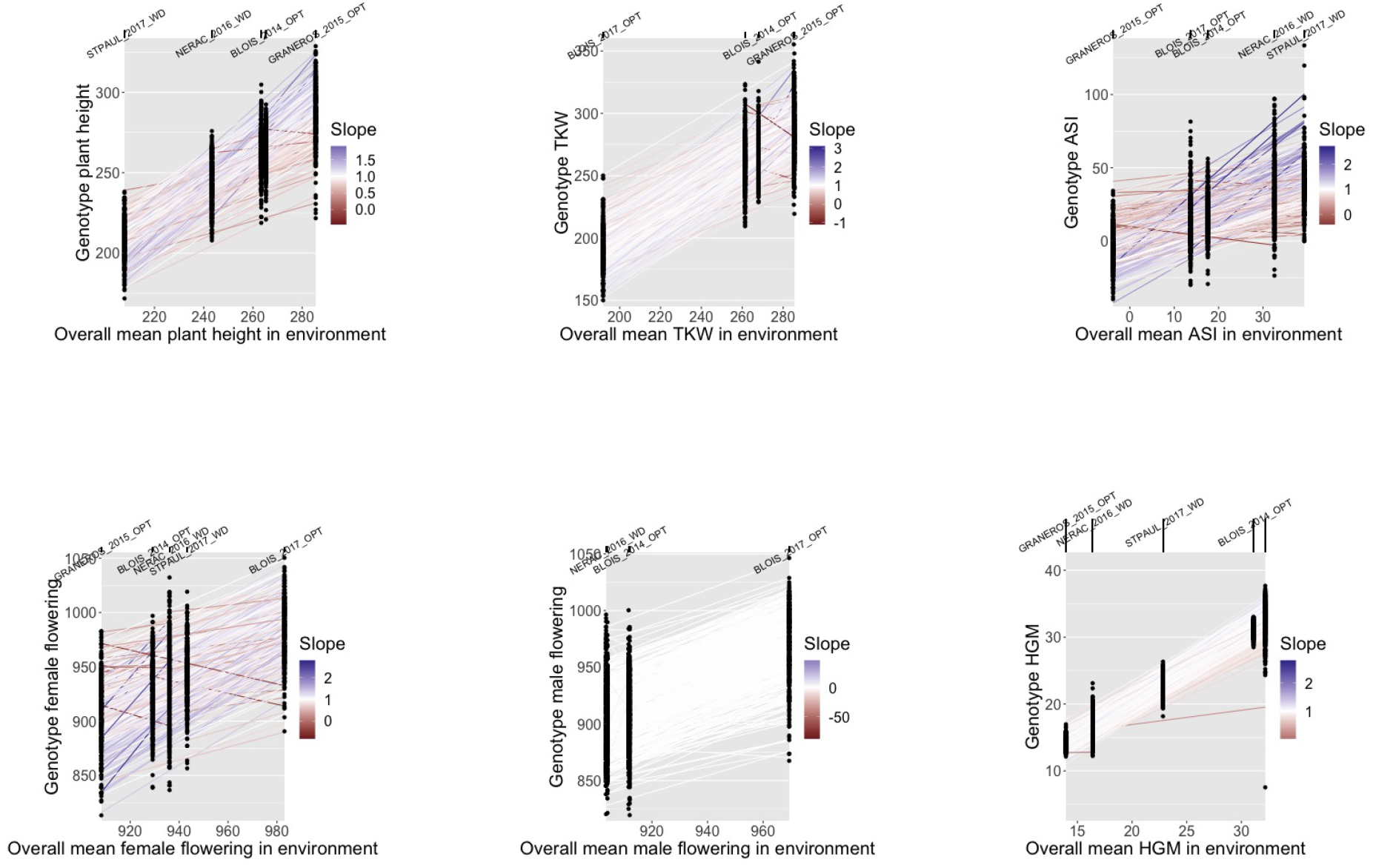
Mean trait values of all genotypes in each environment. On the X axis environments are plotted by the mean of each trait across all genotypes in that environment. Circles are the mean trait values of individual genotypes. Lines are the slope of a genotype’s mean trait value in each environment on the mean trait value of all genotypes in that environment. The color of the line corresponds to the slope; a slope greater (or less) than one indicates a genotype more (or less) responsive to the environment than average.

**Figure S3.**
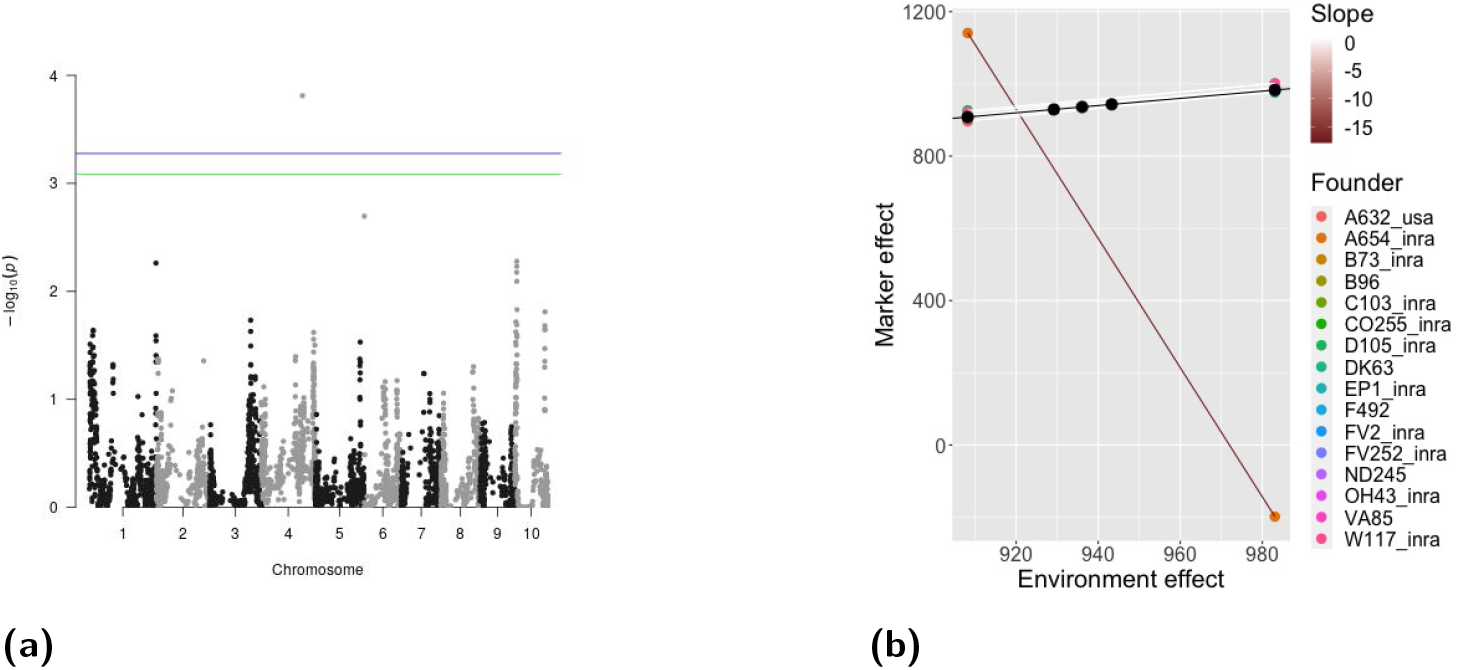
A) Manhattan plot for plasticity GWAS on female flowering. The blue and green lines represent the 5% and 10% significance levels based on permutation tests, respectively. B) Estimated effect of founder ancestry on plasticity for the most significant marker. Lines are the slope of a marker’s effect in each environment on the mean female flowering date of all genotypes in that environment. The color of the line corresponds to the slope; a slope of one indicates a marker with the average response to the environment, a slope less than one indicates a marker less responsive to the environment than average, and a slope greater than one indicates a marker more responsive to the environment than average.

**Figure S4.**
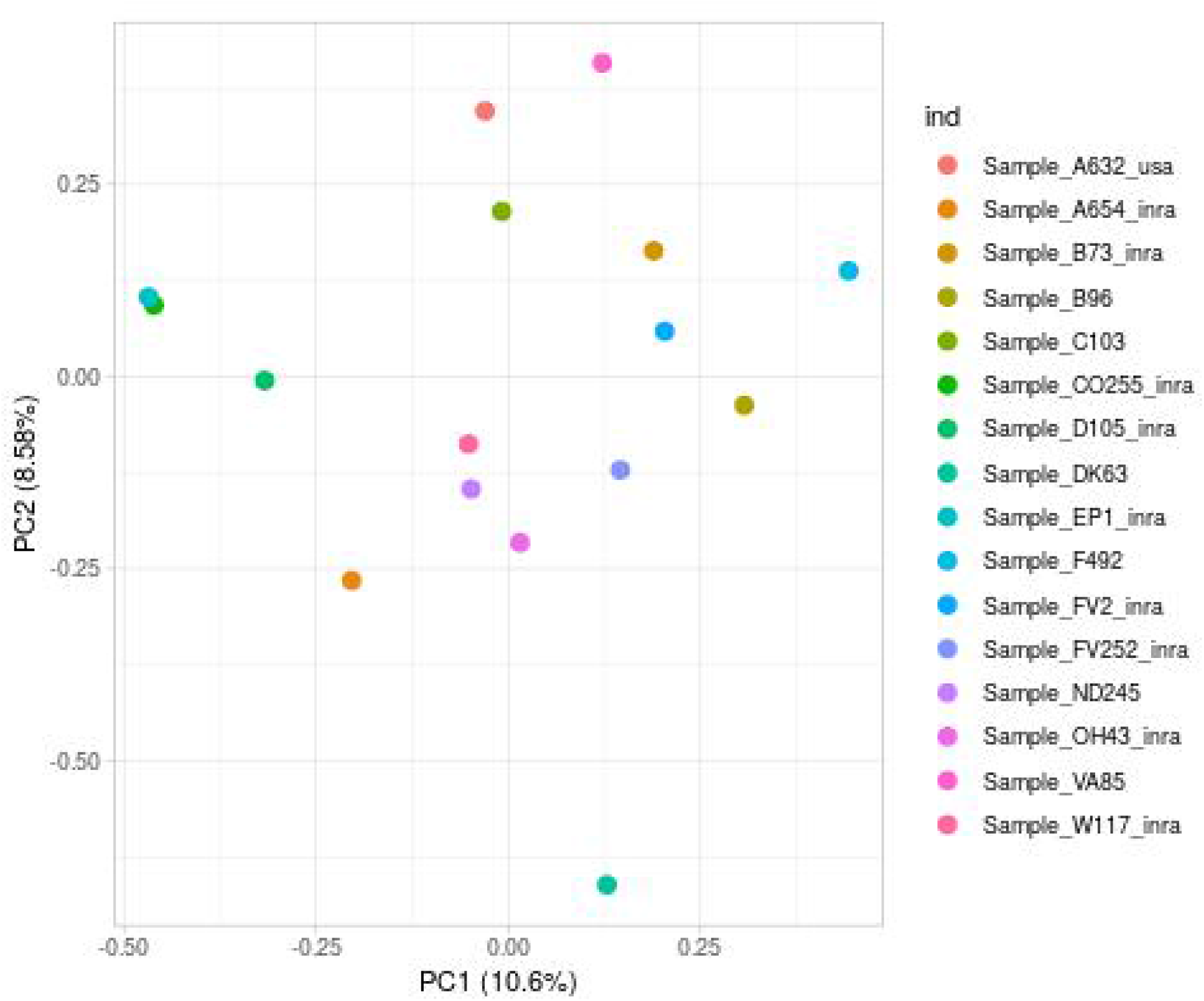
Each founder plotted on the first and second principal components from a principal component analysis of the SNPs within the plasticity GWAS peak for ASI.

**Figure S5.**
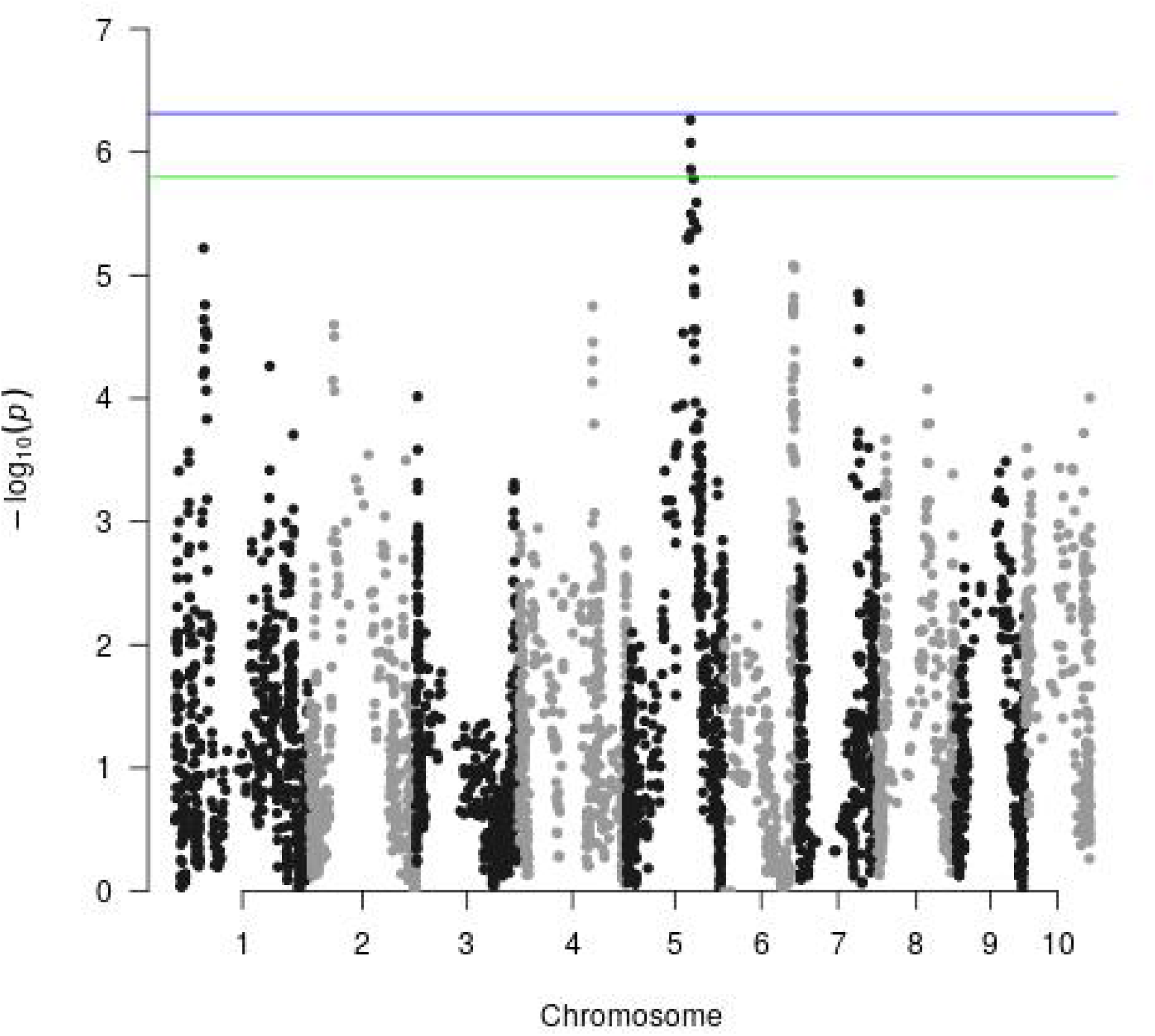
Manhattan plot for plasticity GWAS on grain yield. The blue and green lines represent the 5% and 10% significance levels based on permutation tests, respectively.

**Figure S6.**
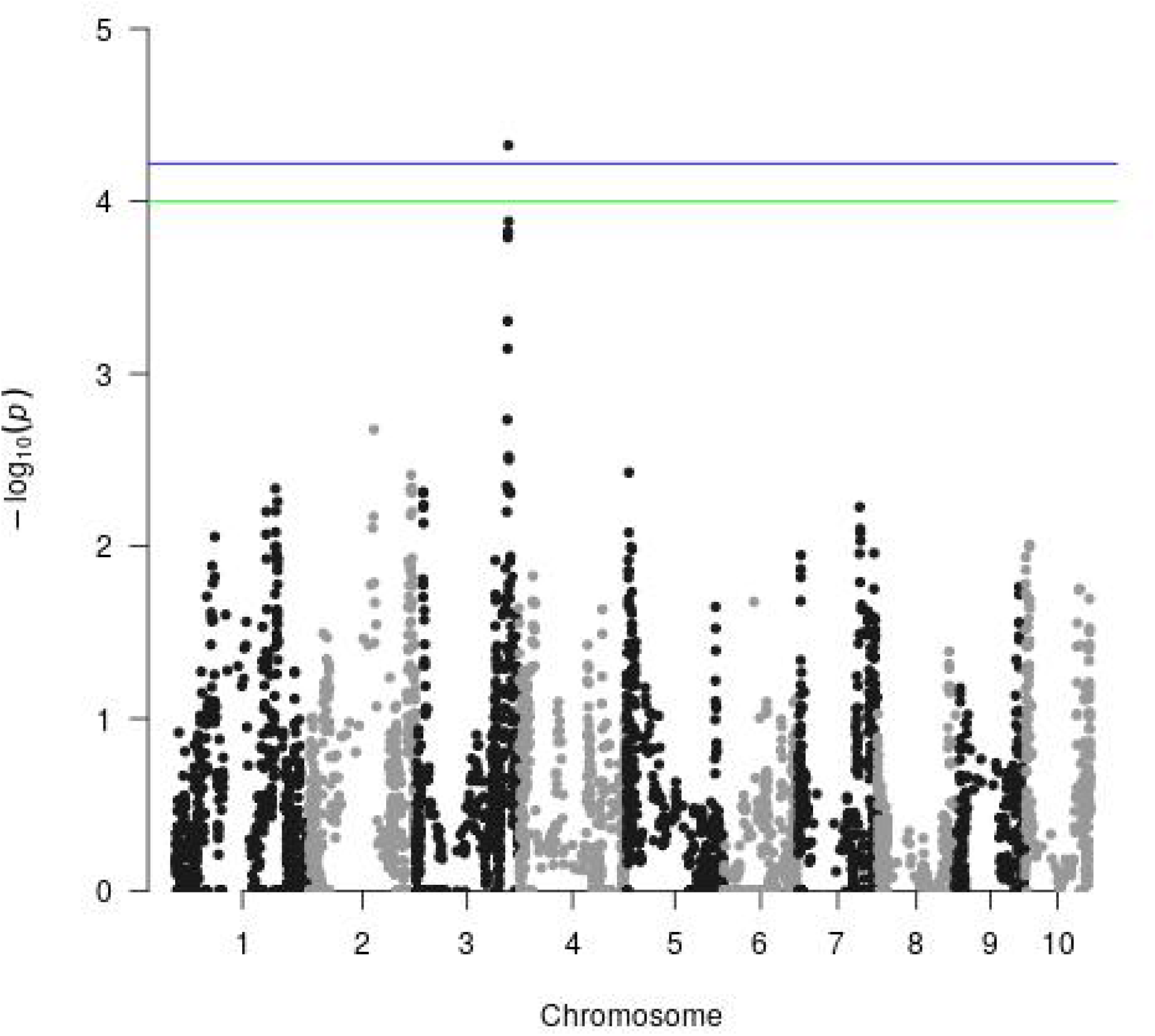
Manhattan plot for Finlay-Wilkinson GWAS on ASI. The blue and green lines represent the 5% and 10% significance levels based on permutation tests, respectively.

**Figure S7.**
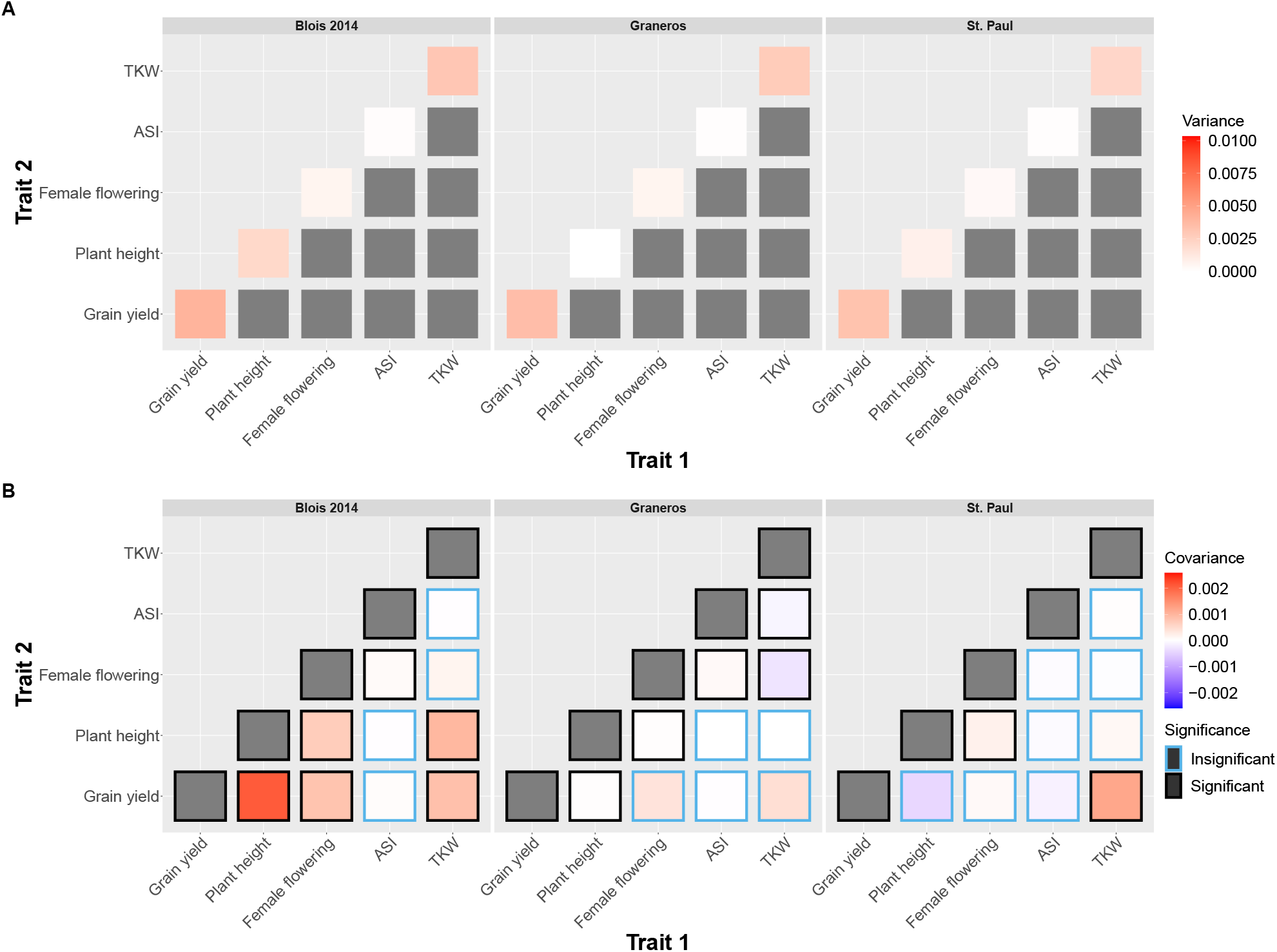
Heat maps of the G-matrices for the remaining environments.

**Figure S8.**
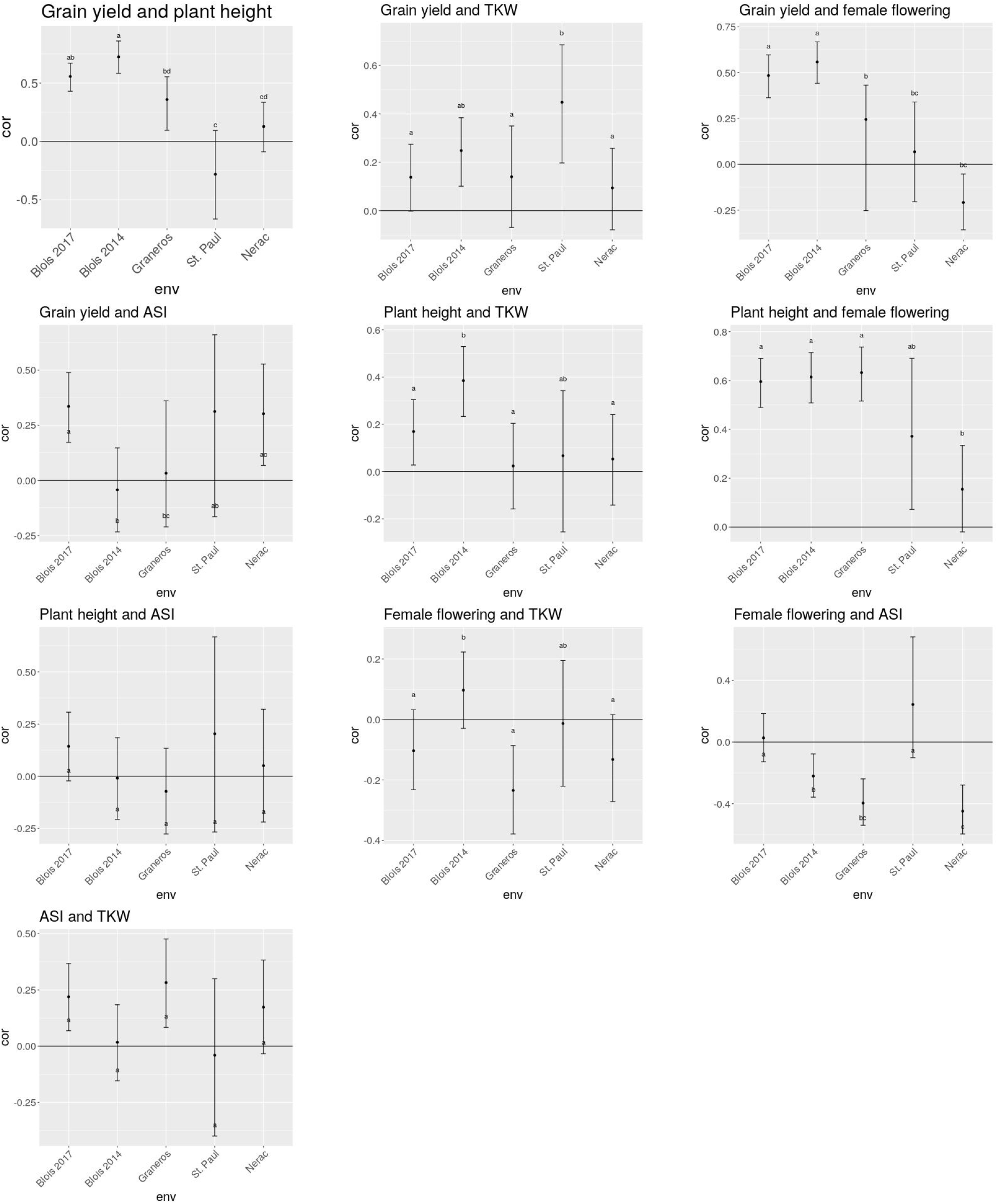
Genetic correlations of each pair of traits. For each pair of traits genetic correlations are shown for each environment with 95% credible intervals. Letters indicate significantly different groups as determined by comparing the 95% credible intervals of the difference between MCMC samples from estimating the correlation in each environment.

**Figure S9.**
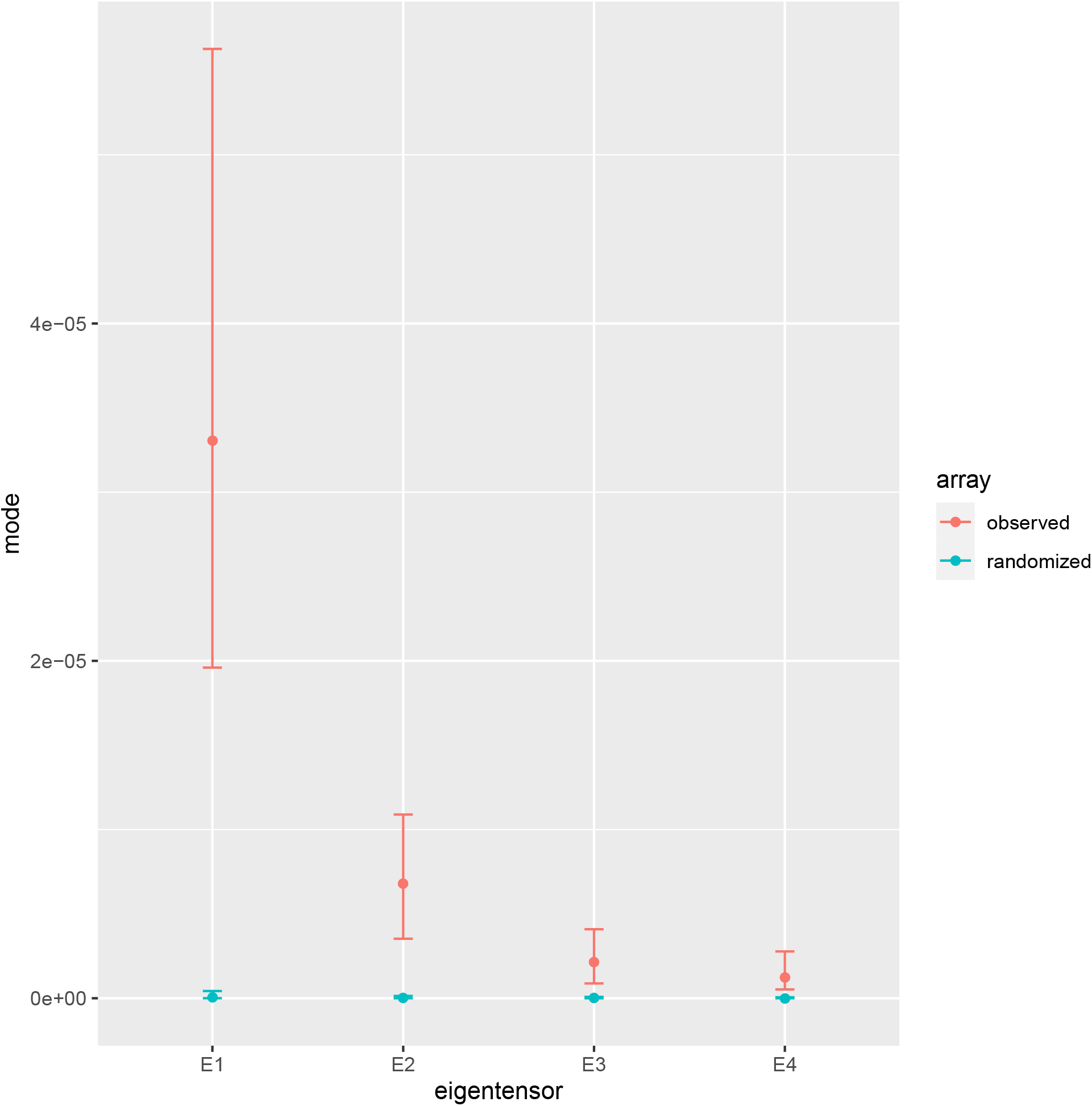
Posterior mode and 95% credible intervals of the eigenvalues of the non-zero eigentensors of the G-matrices. If the observed eigenvalue of an eigentensor is greater than the 95% credible intervals of eigenvalues of the eigentensors estimated from randomized data that indicates the eigentensor explains more of the variation among G-matrices than would be expected by chance.

**Figure S10.**
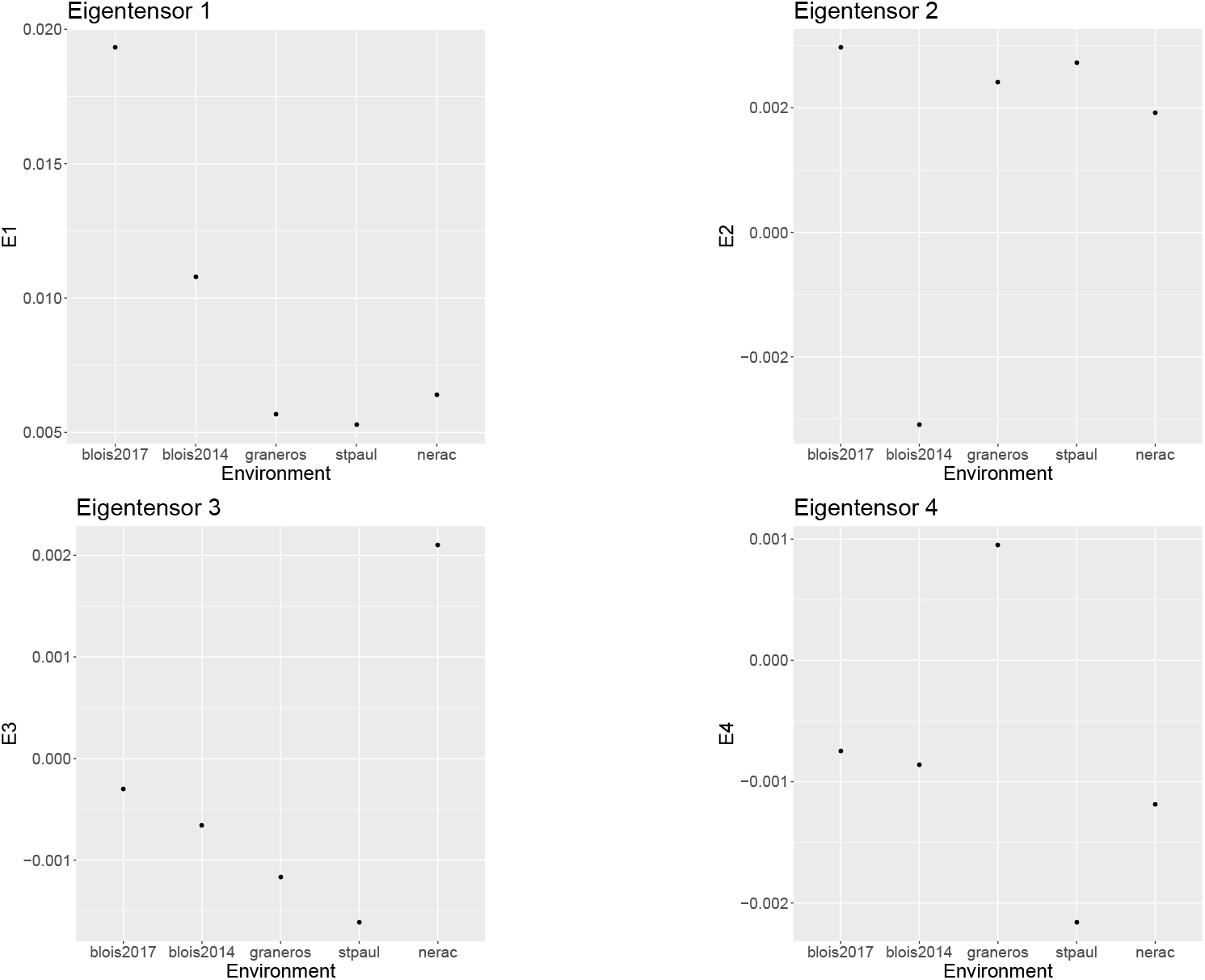
The G-matrix estimated in each environment plotted on each of the four first eigentensors. Note that the scale on the y axis is different for each plot.

